# Communicating Regulatory High Throughput Sequencing Data Using BioCompute Objects

**DOI:** 10.1101/2020.12.07.415059

**Authors:** Charles Hadley S. King, Jonathon Keeney, Nuria Guimera, Souvik Das, Brian Fochtman, Mark O Walderhaug, Sneh Talwar, Janisha Patel, Raja Mazumder, Eric Donaldson

## Abstract

For regulatory submissions of next generation sequencing (NGS) data it is vital for the analysis workflow to be robust, reproducible, and understandable. This project demonstrates that the use of the IEEE 2791-2020 Standard, (BioCompute objects [BCO]) enables complete and concise communication of NGS data analysis results. One arm of a clinical trial was replicated using synthetically generated data made to resemble real biological data. Two separate, independent analyses were then carried out using BCOs as the tool for communication of analysis: one to simulate a pharmaceutical regulatory submission to the FDA, and another to simulate the FDA review. The two results were compared and tabulated for concordance analysis: of the 118 simulated patient samples generated, the final results of 117 (99.15%) were in agreement. This high concordance rate demonstrates the ability of a BCO, when a verification kit is included, to effectively capture and clearly communicate NGS analyses within regulatory submissions. BCO promotes transparency and induces reproducibility, thereby reinforcing trust in the regulatory submission process.

## Introduction

At present, no widely accepted standard exists for reporting NGS (Next generation sequencing) data analysis. This lack of standardization has likely contributed to the “reproducibility crisis” in science^1^. The continued expansion of NGS data and technologies will require researchers, clinicians, industry professionals, and regulatory scientists to collectively address the issues of data reproducibility, standardization, and interoperability^2^. It can be difficult to communicate or interpret big data effectively^3^, as these types of analyses are difficult to share and reproduce. Despite how common data-driven research practices are, good data management is still generally insufficient^4^. Several factors may contribute to the paucity of strong data management practices, most saliently including “inadequacies in the computational and data management approaches available to biomedical researchers. In particular, tools rarely scale to big data^5^.”

Source code is often required to mimic a study; however, very few of the well-known bioinformatics focused journals require authors to include source code and data used in drafting a publication. GigaScience^6^ [(https://academic.oup.com/gigascience/pages/instructions_to_authors]), Elsevier Journal of Proteomics ([https://www.elsevier.com/journals/journal-of-proteomics/1874-3919/guide-for-authors#txt87510]), Genome Research [(https://genome.cshlp.org/site/misc/ifora_overview.xhtml]), ASBMB [(https://www.asbmb.org/journals-news/editorial-policies#data_availability]), and PLOS Computational Biology [(https://journals.plos.org/ploscompbiol/s/materials-and-software-sharing#172])) are a few that request the source code be included when possible. Although requesting source code is a step in the right direction to replicate a bioinformatics analysis pipeline, it is often not enough to understand the study and verify the results.

United States regulatory agencies face an even greater challenge as they typically cannot make a request for something so general. The government guidance needs to be very specific, and contain no legal ambiguities. The US Food and Drug Administration (FDA) has a few regulatory guidance documents pertaining to NGS data that are center-specific^7^; however, none that are applicable agency-wide. In July of 2019, the FDA’s Center for Drug Evaluation and Research (CDER) released a Technical Specification on submitting NGS sequence data^7^. The technical specification was a strong step towards synchronizing communication efforts related to NGS data, and was designed to address questions around how NGS data are generated based on experience with reviews of early NGS submissions: What NGS analysis pipelines are being used? How are these pipelines being validated? What algorithms are being used for specific applications in a pipeline? Is there reproducibility between algorithms? What parameters are used by the pipeline? How are the parameters optimized? How will the analysis pipeline be evaluated for regulatory review? What information will be required to make a regulatory decision? While the technical specification provided a framework for pharmaceutical partners to follow when describing their analyses, it did not specifically address issues related to data reproducibility.

Comparative analysis is essential to effective research and regulatory review so, the lack of submission requirements surrounding *in-silico* or computational experiments in any situation can make it difficult to find all the necessary data, scripts, and algorithms used^8^. The Common Workflow Language (CWL)^9^ is a well-known and established standard, which, as a mechanism for portability of execution, exclusively captures procedural information. Research Objects (ROs)^10, 11^ is another specification that aims to improve the shareability of research and results, and which focuses on operating as an aggregation of all resources needed for execution in a containerized package like Docker ^12^. Both of these specifications adhere to a common set of principles, known as FAIR^13^, or Findable, Accessible, Interoperable and Reusable^14^. CWL is lightweight but only covers procedural practices, while RO has everything included and can be very bloated as a result. A lightweight option to regulate the process of reporting information for a bioinformatics pipeline^15^ is a pressing need.

Early discussions to address the challenge of workflow communication began in 2012 at a small scale, and in 2014, the FDA’s Genomics Working Group^16^ convened a meeting to formally discuss the topic (https://www.fda.gov/media/90504/download). It was determined the resulting solution to workflow communication should satisfy four main criteria:

1. Be human readable.
2. Be computer readable: designed to structure information with predefined fields and associated meanings of values.
3. Contain enough information to understand the computational pipelines, interpret information, maintain records, and reproduce experiments.
4. Be immutable: designed to ensure the information has not been altered.

Funding for a technical solution began shortly thereafter. Five workshops, two publications, and input from over 300 participants associated with academic, government, and private institutions, resulted in the solution now known as BioCompute Object, and which has recently been standardized as IEEE 2791-2020^17^.

BioCompute Object (https://biocomputeobject.org) is a solution that includes a mechanism for recording provenance, metadata, execution environment, and the biological relevance of a computational analysis^14^. Similar to wet lab experiments, a computational analysis may incorporate a multitude of processes, each of which could also require a detailed description of the parameters, inputs, outputs, dependencies, and context for complete understanding of the analysis and its results.

A document that conforms to the IEEE 2791-2020 standard is known as a BioCompute Object (BCO). A BCO abstracts the concept of a computational analysis, making it platform- and tool-independent. Information in a BCO is categorized into conceptually significant “Domains,” making information easy to find. BCOs have been tested in a variety of contexts such as biocuration^18–20^, as a tool for coordinating a large number of geographically separate researchers (in the context of the World Health Organization’s ReSeqTB^21^ efforts); furthermore, as reported in this document, BCOs have been leveraged as a mechanism for transparently communicating the NGS results from a clinical trial to the FDA as part of a regulatory submission.

The work reported here was a collaboration between GW (George Washington University, Washington, D.C., United States), FDA (Food and Drug Administration, United States) and DDL (DDL Diagnostic Laboratory, Rijswijk, The Netherlands) to use a BCO accompanying a regulatory submission: A private industry collaborator (DDL) analyzed data provided by GW using a proprietary pipeline, and submitted the results and a BCO describing the process to the FDA Reviewer, who did not have prior access to the analysis process used by the industry collaborator. This proof-of-concept project began with simulation of NGS data to mimic a clinical trial^22^ submitted it to the FDA. The submission was in the form of a Verification Kit for review as outlined in Figure 1. A Verification Kit consists of the initial data and the accompanying template BCO (tBCO), which can be thought of as the initial blueprint for a more extended analysis. The tBCO can provide the basis for a wide variety of analysis and also used for verification and other researchers can map their own analysis to a tBCO (parent). In contrast with a tBCO, is a run BCO (rBCO). The rBCO can only exist as a derivative of the tBCO, and is a replicate of the tBCO. For example, in this study the DDL Athena tBCO (parent) is the first replicate and it has 99 rBCOs (children) that are derived from the tBCO. The rBCO should only differ from the tBCO (parent) in the inputs and outputs listed.

**Figure 1.**
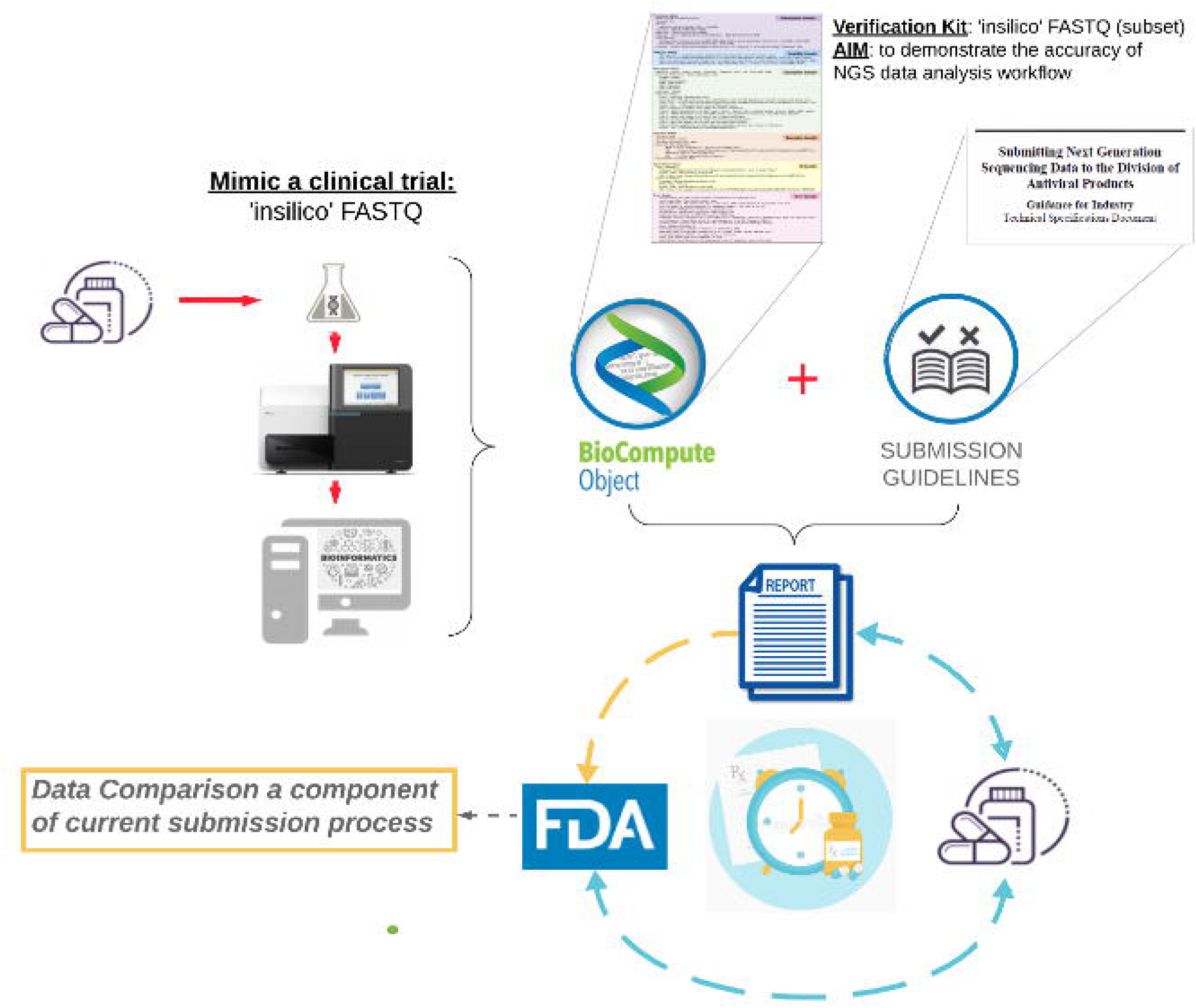
Mock Clinical Trial Next-Gen Sequencing Workflow. To mimic a clinical trial,136 in-silico samples were generated. Of these 36 were representative of a treatment failure population. The data were analyzed and a BCO was generated to report the analysis. The BCO was submitted to the FDA along with the data and a verification kit.

The participant from private industry created a tBCO to report the results of their proprietary pipeline (Figure 2), which had been specifically validated for viral (CMV, HBV, HCV, IAV, IBV, RSV, SARS-CoV-2 and WHV) variant calling analysis of NGS results from patients undergoing viral treatment to assess for resistance-associated amino acid substitutions in the target viral proteins. The DDL Athena pipeline has been previously used to perform variant calling analysis from several clinical trials with different trial phases (1-3). With this project we have specifically mimicked a clinical trial where a combination of drugs against HCV genotype 1a was being tested. Antiviral drugs against hepatitis C virus (HCV) target either nonstructural protein 3/4A (NS3/4A) protease inhibitors, NS5A inhibitors, or NS5B polymerase inhibitors^23^. Treatment type for HCV genotype 1 (GT1) depends on a variety of conditions: subtype, presence of cirrhosis, and considerations of medication and insurance costs (https://www.hepatitisc.uw.edu/go/treatment-infection/treatment-genotype-1/core-concept/all#studies-retreatment-adults-hcv-genotype-1-ledipasvir-sofosbuvir). Different combinations of two or three drugs can clear HCV in more than 90% of patients, including populations that were previously treatment resistant^23^.

**Figure 2.**
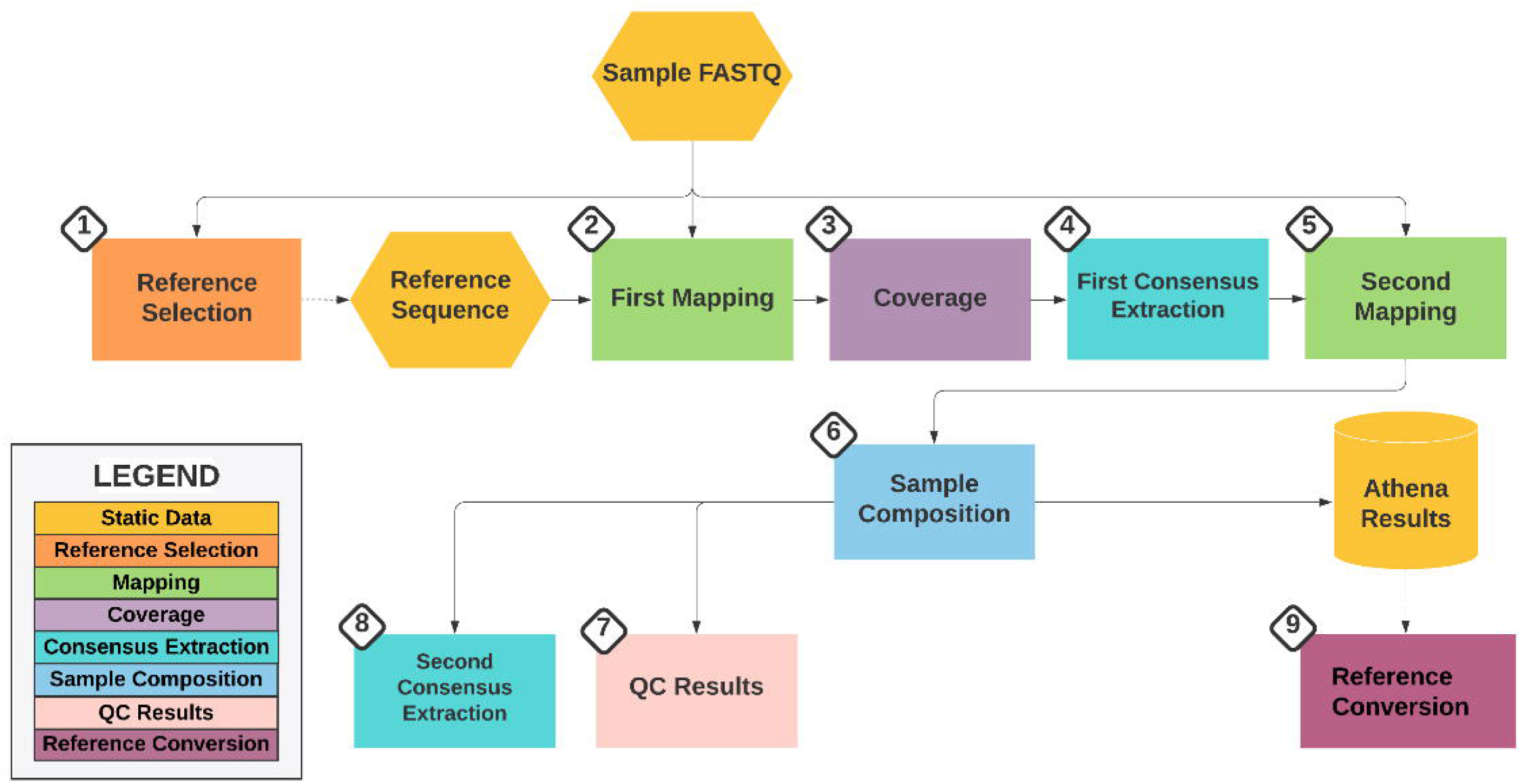
Overview of DDL Athena Pipeline. In the Genotyping/Reference Selection step, 10% of raw reads from *fastq* files are compared against all genotype and subtype reference sequences from a specific organism. After selecting the most appropriate reference, in the First Mapping step, all the raw data reads are mapped against this sequence. In step three, Coverage, the number of reads that reach desired quality threshold on each position are considered, which is further used to construct the consensus sequence. In the First Consensus Extraction step, a consensus sequence per sample, is derived based on coverage values. Insertions and deletions are not built in when creating the first consensus sequence. To improve the mapping statistics and coverage on each position of the reference sequence, in the Second Mapping step, all raw reads are mapped against the first consensus. In step six, Sample Composition, all the differences as well as similarities between the selected official reference sequence and raw reads of a samples are built on amino acid, codon, and nucleotide level. In QC Results step, quality control results are generated which help to determine the quality of the sequencing run of a sample. Using the coverages calculated from second mapping, a second consensus sequence is constructed, in Second Consensus extraction step, which consists of insertions and deletions found in raw data reads. Finally, in the Reference Conversion step, variant records from a genotype or subtype is converted to the universal (main) reference.

Several approved regimens for HCV include sofosbuvir (SOF), which targets the NS5B polymerase inhibitor and inhibits viral replication. Given sofosbuvir is known to have a high barrier to resistance^23^, bioinformatics approaches have been used to “characterize potential resistance-associated substitutions.” A report by FDA collaborators describes the methods to process/analyze NGS data submitted as part of the sofosbuvir (SOF) resistance analysis data set^24^.

To test the utility of a BCO, upon receipt of the Verification Kit (Athena tBCO and the *in-silico* NGS samples), the FDA collaborator did an independent assessment of the reads, generated an FDA tBCO, and then compared the results of the two independent analysis using the tBCO. The goal was to determine: 1) comparability of results; 2) comparability of BCOs; 3) if the FDA can rely upon a tBCO generated by an outside source for accurate representation of the analysis pipeline and the results. The results of both pipelines were in concordance and provided the necessary information about the analysis pipeline. This synthetic clinical trial resistance data set was successfully reproduced using the tBCO as a primary reference in the submission and following the parametric and pipeline steps captured within.

## Results

### Results: Pipeline and example trial

We simulated a 100 patient clinical trial where 18% of the patients experienced a treatment failure after eight weeks, which is tabulated in table 1. The *in-silico* generated samples have amino acid variants at the frequencies indicated in table 2. The variations that were introduced into the reference HCV GT1a sequences, listed in table 3, represented resistance-associated substitutions known to arise against a triple drug combination known as 3D. This treatment regimen includes a protease inhibitor (paritaprevir) + an NS5A inhibitor (ombitasvir) + a polymerase inhibitor (dasabuvir). Here is the scenario: After 8 weeks of treatment, the failure subjects experience a virologic breakthrough (a serum viral load is detected after being undetectable for several weeks) and treatment was stopped for those subjects. The sponsor collected two samples for sequencing, one at baseline prior to the initiation of drug treatment (for all subjects) and one at Week 9 (for treatment failure subjects) when the breakthrough was discovered^25^. The synthetic viral reads were uploaded to the GW public instance of the HIVE platform^26, 27^ and shared with a representative from each of the respective institutions. The full set of reads generated is available for download from https://hive.biochemistry.gwu.edu/datasets#mockHCV1aTrial.

**Table 1.** The relationship between the various simulated patients, samples, and files are tabulated here. We simulated 100 patients with a total of 118 samples which yielded a total of 236 paired-end FASTQ files. Eighteen of the patients were simulated to experience a treatment failure. The treatment failure population had a total of two additional samples taken (4 FASTQ files).

| Items | samples | files |
| --- | --- | --- |
| Patient | 118 | 236 |
| Baseline timepoint | 100 | 200 |
| Treatment failure timepoint | 18 | 36 |
| Items included in verification kit | 2 | 4 |

**Table 2.** The expected substitution frequencies observed in the baseline and treatment failure populations.

| Gene | Substitution (amino acid) | Baseline variation | Treatment failure variation |
| --- | --- | --- | --- |
| NS3 | D168A | 0.05% | 22.00% |
| NS3 | D168Y | 1.10% | 56.00% |
| NS5A | M28T | 1.00% | 71.00% |
| NS5A | M28S | 8.00% | 3.00% |
| NS5A | Q30R | 0.08% | 18.00% |
| NS5B | C316N | 0.00% | 6.00% |
| NS5B | M414T | 3.00% | 44.00% |
| NS5B | S556G | 0.00% | 3.00% |

**Table 3.** The empirical error subdomain represents observed variation or error in a pipeline. This table displays the values of the empirical error domain for both tBCOs discussed in this paper (Athena tBCO and FDA tBCO). The first three columns (Percentage, Reads Generated, and Coverage) were applied for both analyses. **Percentage**: The portion of the entire population that the indicated amino acid substitution represents. **Reads generated**: number mutated of in-silico reads created to obtain the desired coverage and percentage for experiment. **Coverage**: The coverage provided for the genome based on the number of reads generated for the experiment (number of reads generated * read length / genome length). **DDL Athena read count**: number of reads identified by the Athena pipeline with mutation with the indicated variation. **DDL Athena coverage**: Total number of reads mapped to location. DDL **Athena percentage**: Athena determined percentage of identified mutation. **DDL Athena forward count**: Number of reads mapped to the forward strand for this position. **DDL Athena reverse count**: Number of reads mapped to the reverse strand for this position. DDL **Athena FR score**: FR score is a measure of the forward vs revers count for a location. Values < 0.5 are desirable. **HIVE read count**: number of reads identified by the Athena pipeline with mutation with the indicated variation. **HIVE coverage**: Total number of reads mapped to location. **HIVE percentage**: DDL Athena determined percentage of identified mutation. DDL **Athena STDEV.P**: value of STDEV.P for percentage and DDL Athena percentage. **HIVE STDEV.P**: value of STDEV.P for percentage and HIVE percentage.

|  | D168Y | D168A | M28T | M28S | Q30R | C316N | M414T | S556G |
| --- | --- | --- | --- | --- | --- | --- | --- | --- |
| Percentage | 0.56 | 0.22 | 0.71 | 0.03 | 0.18 | 0.06 | 0.44 | 0.03 |
| Reads generated | 90029 | 35369 | 114144 | 4823 | 28938 | 9646 | 70737 | 4823 |
| Coverage | 2800 | 1100 | 3550 | 150 | 900 | 300 | 2200 | 150 |
| DDL Athena read count | 2853 | 1050 | 3652 | 149 | 934 | 297 | 2215 | 127 |
| DDL Athena coverage | 5081 | 5081 | 5158 | 5158 | 5111 | 5002 | 5059 | 4738 |
| DDL Athena percentage | 0.5615 | 0.20665 | 0.70803 | 0.02889 | 0.18274 | 0.05938 | 0.43783 | 0.0268 |
| DDL Athena quality | 33.95 | 33.67 | 33.84 | 33.21 | 33.45 | 33.69 | 33.97 | 32.34 |
| DDL Athena forward count | 1444 | 534 | 1794 | 73 | 449 | 150 | 1103 | 51 |
| DDL Athena reverse count | 1409 | 516 | 1858 | 76 | 485 | 147 | 1112 | 76 |
| DDL Athena FR score | 0.0186 | 0.0284 | 0.004 | 0.0093 | 0.0415 | 0.0466 | 0.0583 | 0.228 |
| HIVE read count | 2853 | 1081 | 3623 | 156 | 947 | 302 | 2206 | 128 |
| HIVE coverage | 5116 | 5116 | 5136 | 5136 | 5077 | 5007 | 5050 | 4739 |
| HIVE percentage | 0.558 | 0.209 | 0.735 | 0.031 | 0.185 | 0.061 | 0.437 | 0.027 |
| DDL Athena STDEV.P | 7.500E-04 | 6.675E-03 | 9.850E-04 | 5.550E-04 | 1.370E-03 | 3.100E-04 | 1.085E-03 | 1.600E-03 |
| HIVE STDEV.P | 1.000E-03 | 5.500E-03 | 1.250E-02 | 5.000E-04 | 2.500E-03 | 5.000E-04 | 1.500E-03 | 0.000E+00 |

### Results: DDL Athena BCO and Verification Kit

The data analysis of synthetic reads by DDL is described in Figure 2. The specific use case of the DDL Athena pipeline verified was for detection of minor HCV GT1a variants on the NS3, NS4A, NS5A and NS5B genes. The objective, to lighten the burden of communication during FDA submission while enabling the discovery of discrepancies found between the data analysis pipelines, was facilitated by using a BCO during our mock clinical trial data submission.

The DDL Athena template BioCompute Object was written collaboratively after the analysis was completed for the first two samples. This exercise resulted in a completed BCO (Supplementary File 1) available at https://w3id.org/biocompute/portal/BCO_00022530 with robust provenance, usability, extension, parametric, input, description domain, and output domains.

The BioCompute Object error domain is used to record values that gauge the accuracy and precision of the analysis the BCO describes. The empirical error subdomain represents observed variation or errors in a pipeline. For this project it was constructed from a tabulated comparison of the known values for the initial baseline sample and the initial treatment failure sample (table 3). This was based on previous work^28^ with the Unified Variant Pipeline (UVP) for ReSecTB^21^, which can be found in the UVP GitHub repository (https://github.com/biocompute-objects/UVP-BCO). The algorithmic error domain represents built-in thresholds for error. The DDL Athena pipeline had a variant detection threshold of 1%, a read quality score requirement of Q>20, and a coverage requirement of 500 reads to call a mutation at that locus. Together with the complete DDL Athena BCO, the two samples constitute the “Verification Kit”, as outlined in Figure 3.

**Figure 3.**
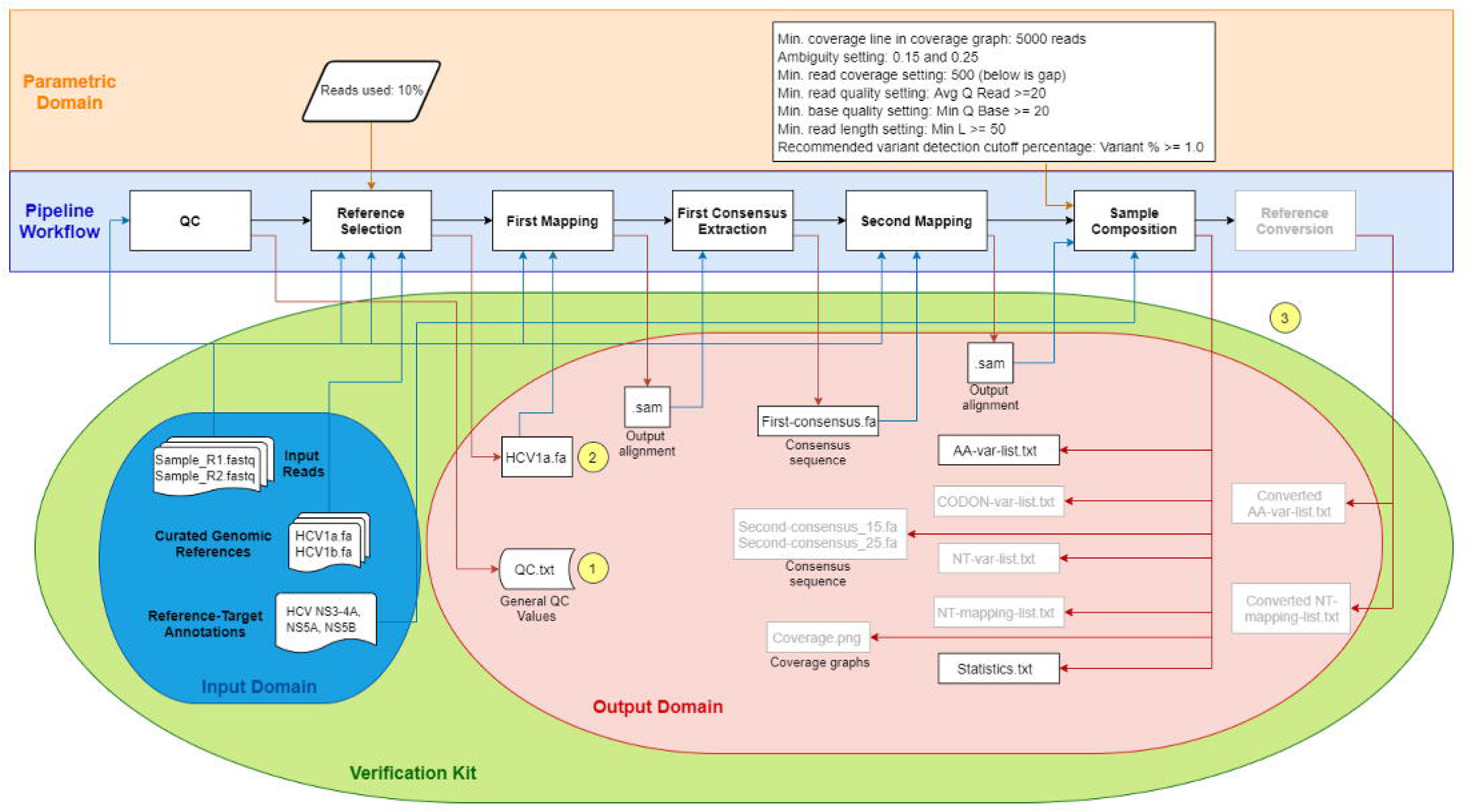
Cartoon Representation of DDL Athena Workflow Highlighting Verification Kit. This figure represents the tBCO from the DDL Athena pipeline. Light blue indicates the pipelines steps described in the description domain: QC, Reference selection, First mapping, First consensus extraction, Second mapping and Sample composition. Reference conversion, although is part of the pipeline, is not highlighted, as this feature is not used in this project. Only the specific pipelines steps used are described in the BCO. The input files listed in the input domain are represented in blue shade. On light red, the output files are represented as listed in the output domain. The different parameters applied on the different pipeline steps are listed in an orange share. In green, the verification kit is represented and specifically determined from the input and output files of this project.

The FDA-CDER reviewers will typically provide the following recommendation for NGS submission to Division of Antiviral products: The Division of Antivirals performs an independent assessment of NGS data for resistance analyses. Please provide the raw fastq files and a frequency table of variations detected along with a detailed description of your NGS analysis pipeline. We encourage you to submit a mock dataset prior to the regular submission to ensure that the information that you intend to submit is acceptable and can be transferred to the agency. For details on what and how to submit NGS data to the division, please see the Guidance for Industry Technical Specifications Document, Submitting Next Generation Sequencing Data to the Division of Antivirals^7^.

The BCO is used to describe the specifics of an NGS data analysis workflow and necessary to determine the quality of the analysis and FAIR principles. As there are so many different analysis methods, many of which are perfectly valid, there is no way to specify all possible workflows.

### Results: FDA reviewer results comparison and verification kit

Samples from all subjects were sequenced at the initiation of treatment (baseline). Eighteen subjects who were simulated to have failed the 3D DAA regimen (paritaprevir+ombitasvir+dasabuvir) to treat their HCV GT1a infection at Week 9 of treatment (treatment failure) had a second sample sequenced at that time point. Supplementary Table 1 lists the 243 comparative variant calls for the entire treatment failure population in the mock trial. Comparison of the two methods show only two of the 242 variant calls (0.8%) were not in agreement. The threshold for agreement between two variant calls required the two following conditions: difference in coverage <500 reads and the difference in variant frequency <0.05%.

This proof-of-concept project demonstrated the utility of using BCO to communicate a complex NGS analysis workflow and the NGS results from that workflow in a robust and easily interpretable format suitable for regulatory review. The BCO provided complete transparency on the NGS data analysis workflow, provided a rational and interpretable structure of the data analysis, contained the exact parameters used for the analyses, and delivered the verification kit from which it was possible to determine that the analysis pipeline was fit for its purpose.

## Discussion

Here, we present the first synthesized trial submission of an analysis recorded as a BCO and submitted to an FDA Reviewer, along with the post-submission Reviewer analysis. This study was designed to simulate the generation of NGS data from a resistance analysis conducted as part of a clinical trial assessing the efficacy of an antiviral drug against HCV GT1a, and the use of BCO to capture the analyses and communicate the results and analysis processes to a regulatory agency. The results of this project indicate that the BCO is capable of capturing the processing workflow of those data by a private entity, and is suitable for submission of the analyses to the FDA. NGS analysis is used to assess a viral target at baseline (prior to drug administration) and near time of failure (on drug treatment) to determine if amino acid changes can be identified that confer resistance to the drug.

Communicating NGS data results has been a persistent challenge. The size of the data and complexity of the analysis often means direct observation is not feasible when evaluating how an analysis was conducted. Other methodologies need to be employed to evaluate the precision and accuracy of an analysis, and many times this is not immediately possible because of missing information. In many instances, an independent assessment of the data submitted for regulatory review has been required because of limitations associated with access to tools & pipeline specifics from the sponsor. Often review and verification of results is difficult because pipeline and parameters are vague, not included, or are treated as a “black-box”. Alternatively, an analysis pipeline may contain manual data analysis that is not described in great detail, if at all. All of these scenarios are examples of common but avoidable complications in the current ad hoc system of method communication, and underscore the need for more thorough communication and clarification.

Both FDA and DDL received the reads, did an independent analysis, created a tBCO, generated the subsequent rBCOs, and aggregated the results. Both groups compared the results and agreed that they were in concordance. The FDA reviewer compared tBCOs and concluded that the DDL BCO and the FDA BCO were comparable, that the DDL tBCO was interpretable, and that it provided sufficient information to perform a regulatory review of the NGS data.

Using a tBCO also enabled us to address each of the questions outlined in the introduction. *How are the data being analyzed?* The specific variables used to filter the results and conduct quality assurances were all listed. *Are the results robust?* By comparing the DDL and FDA analysis we demonstrated that the results were robust, despite being generated on different analysis pipelines. And in turn each BCO for the respective pipeline demonstrated the robustness of the results via the error domain. *Are the results reproducible?* The use of DDL’s Athena BCO to create the HIVE BCO demonstrates that the results are reproducible, and that the BCO mechanism is platform independent. *What programs and parameters are used?* Each of the tools used is recorded with the inputs, outputs, dependencies, and parameters. *Is the analysis pipeline publicly available?* While the public availability of all aspects of a pipeline is not required to use or compose a valid BCO, the methods presented here can be easily applied in an open data context.

The BCO is used to describe the specific details of this NGS data analysis workflow that are missing on the current guidance, but that are necessary to determine the quality of the analysis and adherence to FAIR principles. As there are so many different analysis methods, many of which are perfectly valid, there is no way to specify all possible workflows.

During regulatory review the questions a reviewer must answer are “What data are necessary to make a regulatory decision?”, “Are summary data from one analysis pipeline sufficient?”, and “How will the analysis pipeline be validated?”. The BCO (IEEE-2791-2020) standard ensures that the content to answer these questions are included, and the inclusion of a verification kit answers these questions explicitly. The use of a full Verification Kit provides the scaffold needed for a workflow/pipeline to adhere to FAIR data and reproducibility principles.

## Methods

### Methods summary

To provide strong evidence for a proof-of-concept, it was critical to replicate an actual clinical trial in as much detail as possible. The sequencing files reported by a published HCV clinical trial (https://clinicaltrials.gov/ct2/show/NCT02613403) were reproduced in the *in-silico* samples we created. The amino acid substitutions outlined in table 2 was used to create *in-silico* reads which were distributed to DDL for analysis, and represent the full scope of the analysis in a BCO. Once the analysis was completed, the samples were transferred to the FDA reviewer via HIVE along with the BCO. No other information was provided to the reviewer. The FDA reviewer then recreated the analysis using HIVE tools and compared the results. The generalized workflow is outlined in Figure 1.

### Analysis design and example trial

To replicate the example clinical trial, the following scenario was developed: A population of individuals (n=100) chronically infected with Hepatitis C Virus are treated with a triple drug combination known as 3D, which includes a protease (NS3-4A) inhibitor (paritaprevir) + an NS5A inhibitor (ombitasvir) + a polymerase (NS5B) inhibitor (dasabuvir). The sponsor collected two samples for NGS, one at baseline prior to the initiation of drug treatment and one at Week 9, to follow up for identification of drug resistance substitutions in subjects who failed treatment. We mimicked the presence of 18% (n=17/94) samples with a viral breakthrough at Week 8 as indicates a treatment failure and implies the stop of treatment. A viral breakthrough is analyzed in this study by comparing the amino acid % detected by NGS at baseline compared to sequences derived from Week 9 samples. The Week 8 sample is considered to be the timepoint when resistance would be observed based on predetermined sampling and the Week 9 sequencing represents when the sample was sequenced.

The simulated sequencing was designed to replicate 150 base pair Illumina MiSeq, with an output of paired end reads in FASTQ format.

### Synthetic read generation for trial and verification kit

Given the lack of intronic regions in viral genomes, the first base of the codon **(C1)** corresponding to a given protein residue (R) could be calculated using the formula

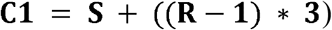

where S is the position in the genome where the gene starts. For each desired mutation the protein residues corresponding to the codon of interest as well as for two codons before and after were confirmed to match the reference (table 4). The nucleic acid mutation applied to the *in-silico* sequence was the one that generated the desired protein substitution with the fewest number of nucleic acid changes.

**Table 4.**
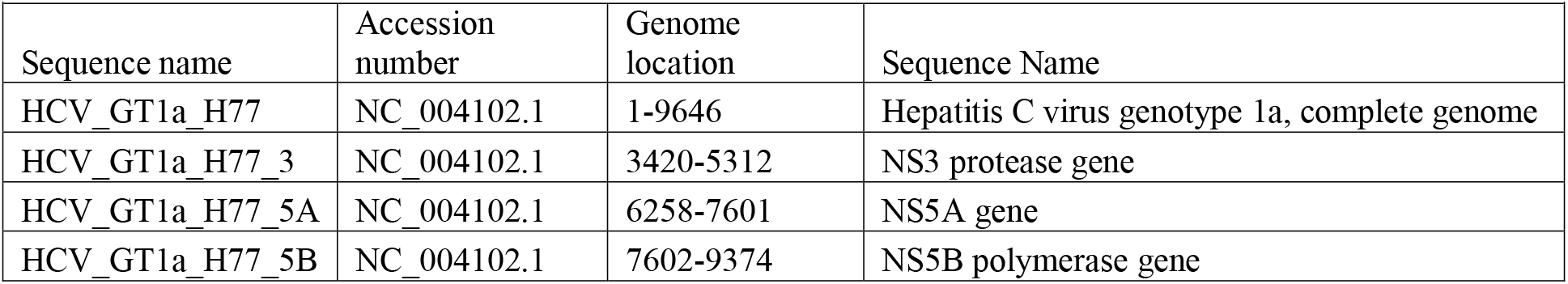
Sequences used to generate the in-silico reads for the paired-end baseline samples (n=100) and for the paired-end Treatment Failure samples (n=18).

| Sequence name | Accession number | Genome location | Sequence Name |
| --- | --- | --- | --- |
| HCV_GT1a_H77 | NC_004102.1 | 1-9646 | Hepatitis C virus genotype 1a, complete genome |
| HCV_GT1a_H77_3 | NC_004102.1 | 3420-5312 | NS3 protease gene |
| HCV_GT1a_H77_5A | NC_004102.1 | 6258-7601 | NS5A gene |
| HCV_GT1a_H77_5B | NC_004102.1 | 7602-9374 | NS5B polymerase gene |

Single codon substitution genomes were created as described above for all given protein substitutions. For the ‘Baseline sample’ read set, the sum of the populations containing any substitution was less than 100%. Reads could therefore be generated by combining reads from mutated genomes with single mutations (supplementary table 2). The number of reads generated by the HIVE DNA-InSilico tool^26^ for each variant sub-population was directly proportional to the protein substitution within the entire population, i.e., for a substitution with a frequency of 8%, eight reads were generated. The remaining reads were replicated directly from the wild type (reference) genome. Reads were then divided between the 100 pseudo-sample read pair files with care taken to preserve order so as to retain the utility of the paired end reads.

The *‘Treatment Failure sample’* reads required the generation of multi codon substitution genomes because the sum of protein substitutions in the *in-silico* population was greater than 100%. This added a significant level of complexity to the read generation process. Multi codon substitution genomes were created as above combining nucleic acid mutations at all nonconflicting positions. For each mutation a number of reads was determined proportional to the given percentage of the mutant within the *in-silico* population, this added to higher than the number of reads needed to create the coverage of interest. Read creation followed an iterative process. First, a multi mutated genome containing all nonconflicting mutations was created. HIVE DNA-InSilico was used to create enough reads from this genome to cover the number of reads needed for the rarest desired substitution. Second, the number of reads needed for all substitutions present in this multi-mutated genome were reduced by the number of reads created. These steps were repeated until no more mutant reads were needed and the remaining coverage was obtained from the wild type reference genome (supplementary table 3). Reads were divided as for the ‘baseline’ pseudo-sample. The synthetic viral reads were uploaded to the GW public instance of the HIVE platform and shared with a representative from each of the respective institutions. The full set of reads generated is available for download from https://hive.biochemistry.gwu.edu/datasets#mockHCV1aTrial.

### Clinical trial data analysis by DDL Athena Pipeline

The DDL Athena virology pipeline (https://www.ddl.nl/bio-informatics/#athena-virology-pipeline) was used to analyze the in-silico NGS reads created to mimic the HCV clinical trial. This variant calling analysis pipeline has been specifically developed and validated, as required for clinical trials evaluating antiviral drugs, where viral resistance substitutions should be monitored.

The flowchart (Figure 2) shows the different steps of the DDL Athena-pipeline. The initial analysis of DDL Athena is to perform Quality Control (QC) on the input dataset. QC values are determined (number of reads, average read length, % of reads with average Q≥30), which can indicate bad data quality. After QC analysis, reference selection is carried out, using 10% of reads, to determine the specific reference to be used on the first mapping step, by assigning each of the reads to one of the selected references, using Bowtie2^29^ and storing the aggregate results in an SQL database. In the first mapping step, reads are aligned to the selected reference using Bowtie2^29^. The coverage is calculated, by calculating the per nucleotide (A, T, G, C and DEL) per position read counts and, subsequently, a first consensus sequence is extracted for each sample. In the second mapping step, the sample reads are remapped to the extracted first consensus sequence to increase the accuracy of the mapping and obtain maximum coverage. Finally, in the sample composition step of the DDL Athena pipeline, the position of the reads in the second mapping is used to compare the read contents with the original reference sequence. From this comparison, similarities as well as differences (substitutions, deletions and insertions) between reads and the reference sequence can be identified. This comparison is performed on nucleotide, codon and amino acid levels to ensure the traceability of the analysis and to allow viral sequence variation on these different levels. If data analysis is required to compare the variant calling results between different viral genotypes (e.g. HCV1, HCV2 and HCV3), DDL Athena can translate the individual genotype results to an universal reference (e.g. HCV1) and report the results accordingly. As seen in Figure 3 the specific features of the DDL Athena analysis applied in this study are highlighted and the different parameters used are listed. On completion of an Athena analysis different files can be reported: sample consensus sequence *(fasta* files) (similar to Sanger sequencing) with different ambiguity percentage if required (e.g. 25% or 15%), tab-delimited text files consisting of variant data on all aforementioned levels (NT mapping level and Target sublevels) that include all the reference and variant records as derived from the read mappings along with record statistics, the gap tables with regions treated as gaps based on the minimum coverage threshold, a summary of the coverage achieved for each target, a full genome coverage graph and the per target coverage graphs (png files). Finally, also a high-level overview of this data is presented in an automatically generated pdf report for every sample. DDL Athena reports are aligned with the latest FDA recommendations on how to report next generation sequencing data.

During this study the specific data analysis performed by DDL Athena have been described in the provenance, usability, description, execution, parametric, input and output domains. The error domain was not included in the initial proof of concept but developed after the analysis had been completed. The finished BCO is included in 1.

When creating the Error Domain and verification kit to represent the DDL Athena analysis, the strategy built on previous work^28^ with the Unified Variant Pipeline (UVP) for ReSecTB^21^, which can be found in the GitHub repository for UVP (https://github.com/biocompute-objects/UVP-BCO).

The variant calling results were tabulated (Table 3) against the known variant proportions of the initial baseline sample and the initial treatment failure sample. The standard deviation of the population between the known variant percentages and called variant percentages was used to assess the actual error. Column descriptions are below, and are summarized in the table. The table was then converted to a JSON object with each row as an object.

Given a reference length of 9,646, a desired coverage of 5,000, and a read length of 300 (150 b.p. paired end reads), the total number of reads was calculated as (reference length • coverage / read length), or 160,767 reads. Synthetically introduced mutations were generated sequentially, according to the table.

“**Percentage**”: Proportions of variants desired in each sample (baseline and Week 9), expressed as a percentage of reads, taken from the representative clinical trial.

“**Reads Generated**”: The number of reads that sequenced the mutation in question. The value was calculated as the total number of reads for that experiment (calculated using the Lander/Waterman equation[PMID: 3294162]), multiplied by the percentage of reads representing the mutation.

“**Coverage**”: Mean coverage per read times the number of reads representing that mutation.

“**Mutation Call Prob**”: Probability that the DDL Athena pipeline will call the mutation in question. The Poisson distribution was used to calculate the probability that a mutation call would be made for each sample, based on the given parameters. For a population of genomes, the expected rate λ was calculated as total coverage • the proportion of the sample containing the mutation. The DDL Athena pipeline will make a mutation call if >= 1% of the reads representing a particular locus contain a variant, or λ • 1%, and this was used as the number of occurrences *k.* The probability was calculated as 1 - the cumulative probability of 0 through n-1, where n is the detection threshold.

The intended use of the DDL Athena pipeline to be verified was the detection of minor HCV variants on NS3, NS4a, NS5A and NS5B. After FDA data submission, the data set was reanalyzed by the FDA pipeline, allowing for the detection of discrepancies between pipelines during data revision. The objective was to determine if the BCO facilitates FDA submission and to investigate possible discrepancies found between data analysis pipelines.

### FDA reviewer generated BCO and results comparison

The DDL Athena BCO was uploaded to HIVE when completed by DDL and shared with the FDA reviewer. To more faithfully recapitulate an actual regulatory submission, the FDA

Reviewer did not have access to the data and procedural steps prior to submission. GW participants acted as intermediaries to keep information blinded. Using the BCO and synthetic reads generated, a workflow was constructed using HIVE-hexagon^30^ and HIVE heptagon ^26, 27^. A BCO was created using the HIVE BCO tools and is included as #HIVE_BCO.json.

The treatment failure samples of the synthetic reads were all reanalyzed using the HIVE pipeline described above. Data quality control was also done by HIVE. Variant calling was done using the HCV1a genome and the variant calls for each gene (NS5a, NS3, NS5b) from the HIVE pipeline were output to a CSV and combined with the DDL Athena variant calls to determine if the two methods were in “agreement” (Supplementary #table AnalysisComparison). The agreement was determined by the following conditions:

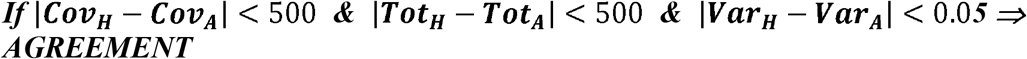

where *Cov_H_* is the variant coverage detected by HIVE, *Cov_A_* is the variant coverage detected by DDL Athena, *Tot_H_* is the total coverage detected by HIVE, *Tot_A_* is the total coverage detected by DDL Athena, *Var_H_* is the variation percentage calculated by HIVE, and *Var_A_* is the variation percentage calculated by DDL Athena. This set of conditions requires that the two methods have a variant coverage within 500 reads of each other and a frequency within 0.05. Given these conditions only two of the 242 variant calls (0.8%) were not in agreement.

The empirical error subdomain for the HIVE BCO was tabulated the same way as the DDL Athena BCO empirical error subdomain. Table 3 was converted to the JSON format and included as the empirical error subdomain in the HIVE BCO. The algorithmic subdomain was populated using the variant frequency cutoff value of 1% and the quality score cutoff of 25.

## Supplementary files legends

Supplementary File 1: Athena.json

Supplementary File #HIVE.json

Supplementary Table 1. The comparative variant calls for all 236 samples in the mock trial submission. Based on a set of conditions determined by Donaldson it was determined that the two methods only two of the 242 variant calls (0.8%) were not in agreement. The conditions required that to be “in agreement” each variant call have a variant coverage within 500 reads of each other and a frequency with in 0.05%.

Supplementary table 2. The number of reads generated by the HIVE DNA-InSilico tool for each mutation was directly proportional to the population of the corresponding protein mutation within the population. The remaining reads were taken from the wild type reference genome. Reads were then divided between the 200 pseudo-sample read pair files with care taken to preserve order in order to retain the utility of the paired end reads.

Supplementary table 3. These reads required the generation of multi codon substitution genomes because the sum of protein mutations in the *in-silico* population was greater than 100%. This added a significant level of complexity to the read generation process. Multi codon substitution genomes were created as above combining nucleic acid mutations at all nonconflicting positions. For each mutation a number of reads was determined proportional to the given percentage of the mutant within the *in-silico* population, this added to higher than the number of reads needed to create the coverage of interest. Read creation followed an iterative process. First, a multi mutated genome containing all nonconflicting mutations was created. HIVE DNA-InSilico was used to create enough reads from this genome to cover the number of reads needed for the rarest desired mutation. Second, the number of reads needed for all mutations present in this multi-mutated genome were reduced by the number of reads created. These steps were repeated until no more mutant reads were needed and the remaining coverage was obtained from the wild type reference genome.

